# DeepAllo: Allosteric Site Prediction using Protein Language Model (pLM) with Multitask Learning

**DOI:** 10.1101/2024.10.09.617427

**Authors:** Moaaz Khokhar, Ozlem Keskin, Attila Gursoy

## Abstract

Allostery, the process by which binding at one site perturbs a distant site, is being rendered as a key focus in the field of drug development with its substantial impact on protein function. The identification of allosteric pockets (sites) is a challenging task and several techniques have been developed, including Machine Learning (ML) to predict allosteric pockets that utilize both static and pocket features. Our work, DeepAllo, is the first study that combines fine-tuned protein language model (pLM) with FPocket features and shows an increase in prediction performance of allosteric sites over previous studies. The pLM model was fine-tuned on Allosteric Dataset (ASD) in Multitask Learning (MTL) setting and was further used as a feature extractor to train XGBoost and AutoML models. The best model predicts allosteric pockets with 89.66% F1 score and 90.5% of allosteric pockets in the top 3 positions, outperforming previous results. A case study has been performed on proteins with known allosteric pockets, which shows the proof of our approach. Moreover, an effort was made to explain the pLM by visualizing its attention mechanism among allosteric and non-allosteric residues.

## 1. Introduction

Allostery is a mechanism that regulates protein activity through a ligand binding at a distant site that is different than the active site. Most drugs change the activity of a protein by directly binding to the active pocket. It has been suggested that every protein possesses allosteric behavior. Even if a certain protein has not yet exhibited allosteric behavior, it could be because of the absence of right conditions such as allosteric effectors or certain mutations (Gunasekaran et al., 2004; Tsai et al., 2008). Allosteric drugs offer the advantage of having fewer side effects than that of orthosteric drugs (Mannes et al., 2022). In contrast to allosteric sites, active sites are highly conserved across a protein family; a drug may bind to the active site of several members of a family. Moreover, allosteric drugs bind elsewhere on the protein surface: surface regions are less conserved across families and it gives us the benefit that when specialized drugs are difficult to bind, effective allosteric drugs can be made (Nussinov and Tsai, 2012).

Several ML methods use pocket features to predict allosteric pockets (sites) (Huang et al., 2013; Greener and Sternberg, 2015; Song et al., 2017; Bian et al., 2019). Notable related works are PASSer, PASSer 2.0, and PASSerRank (Tian et al., 2021; Xiao et al., 2022; Tian et al., 2023b) that follow the similar approach by extracting pockets through FPocket (Guilloux et al., 2009). FPocket, gives feature vectors representing each pocket and further they train several models on these extracted features by performing binary classification i.e. whether a given pocket is allosteric (positive) or not (negative). However, they have not leveraged the power of pre-trained Protein Language Models (pLMs) or Protein Large Language Models (pLLMs). We fine-tuned ProtBERT-BFD (ProtBERT-Big Fantastic Database) pLM from the family of ProtTrans (Elnaggar et al., 2021) on the AlloSteric Database (ASD) dataset (He et al., 2023) and extended it by fine-tuning the pLM in Multi Task Learning (MTL) fashion/setting by using two prediction heads that predicted (A) allosteric residues (B) secondary structure residues. Task A is the primary task and the idea was that in addition to unavailability of large allosteric dataset, while learning allosteric residues (tokens), the model could get information from the secondary structure of the protein and hence would get better information in order to learn allosteric residue features. Further, we leveraged these fine-tuned pre-trained Language Models (pLMs) as backbones (feature extractors) by combining their features with FPocket features to train XGBoost and Automated Machine Learning (AutoML) models. These pLM features’ based trained model outperform the performance of already available computational approaches and prove that the pLM features do provide useful information that helped us achieving better performance scores. To support our concept, we provide the internal attention mechanism to explain the fine-tuned pLM and a case study that shows correctly predicted allosteric pocket.

## 2. The Approach

### 2.1. Dataset

AlloSteric Database (ASD)^1^ is an annually updated, freely available collection of allosteric proteins (He et al., 2023). We followed similar preprocessing steps as described by Tian et al. (2023b), adapting the provided Python scripts^2^ to clean the dataset. These steps involved selecting protein structures with a resolution below 3 Å, ensuring completeness (i.e., no missing residues), and maintaining a sequence identity of less than 30%. Additionally, MMseqs2 (Steinegger and Söding, 2018) was used to filter out proteins with sequence similarities exceeding 30%. MMseqs2 clusters sequences using their similarity hence a representative protein was selected from each cluster. After preprocessing, a total of 207 proteins were extracted from the source ASD dataset and randomly split into 80% for training and 20% for testing. Table 1 gives the detail of the split. A full list of dataset is given in the supplementary material (S1).

**Table 1.**
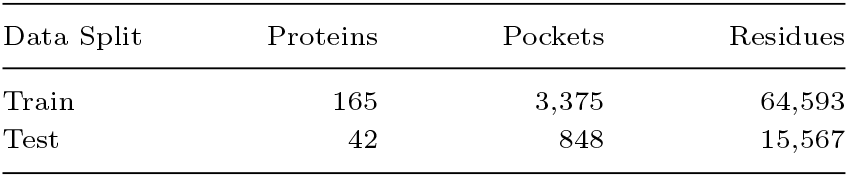
Train/Test Split.

| Data Split | Proteins | Pockets | Residues |
| --- | --- | --- | --- |
| Train | 165 | 3,375 | 64,593 |
| Test | 42 | 848 | 15,567 |

On average, ~20 pockets were detected in each distinct protein. A huge class imbalance was detected where positive samples (pockets) accounted to only 304 or 7.76% of the dataset (4223 pockets). Moreover, at the residue level, there were 5.12% positive labels in the dataset.

### 2.2. Methodology

Figure 1 gives the overview whole architecture of our study. A protein structure and sequence is fed into FPocket and finetuned ProtBERT pLM, respectively. FPocket extracts pockets, where each pocket has PDB file format like coordinates and a 19-*d* feature vector. The pLM also produces features where each vector is of size 1024 and represents a single residue in the sequence. All 1024-*d* vectors representing residues in a certain pocket are aggregated (averaged) to make one 1024-*d* vector. Both feature vectors (from FPocket and pLM) are concatenated, resulting in a 1043-*d* vector. This feature vector is further fed into XGBoost and AutoML models, representing a pocket, and these models classify whether this pocket is an allosteric pocket or not.

**Fig. 1.**
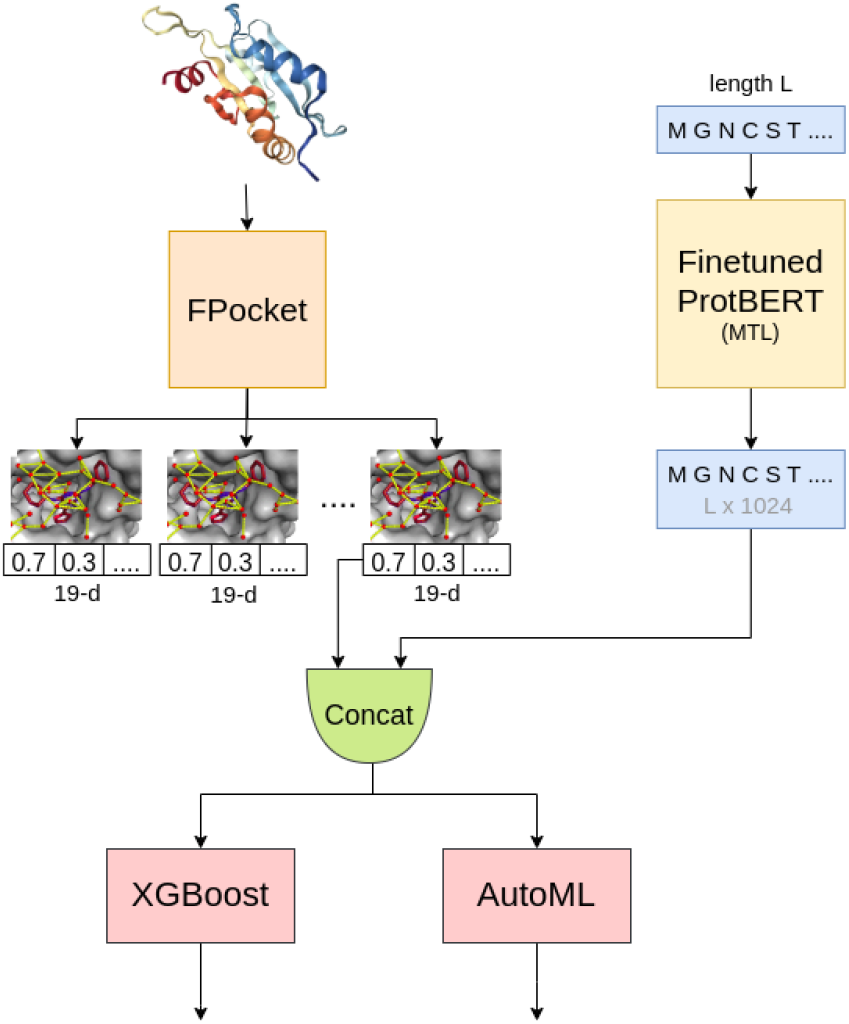
Architecture and Approach. FPocket extracts pockets represented by a 19-*d* feature vector and the finetuned pLM gives feature vectors of size 1024. Feature vectors (from the pLM) representing residues inside a certain pocket are aggregated to produce a single 1024-*d* feature vector and concatenated with a 19-*d* FPocket feature vector resulting in 1043-*d* vector (input to the AutoML and XGBoost models).

The approach of finetuning and leveraging pLMs, from our knowledge, is novel for allosteric pockets prediction and has never been used before.

#### 2.2.1. pLM Finetuning Setup

Elnaggar et al. (2019) implemented MTL to train a pLM on different tasks i.e. to predict secondary structure (SS) and function of protein. By fusing multiple tasks, each task got information from other tasks that helped a certain task to predict better. Following this idea, we prepared the model to get structure level information in combination with other tasks in order to improve the model’s performance to predict allosteric residues (as primary task).

The pLM was fine-tuned to perform residue-level prediction at each head. First, to feed the allosteric dataset into the model, following steps were performed:

- Each residue in an allosteric pocket was labelled as allosteric. Hence, each residue in a protein sequence was labelled as either positive or negative referring to being allosteric or non-allosteric, respectively.
- Each residue in each sequence was separated by *space* character, as the expected input of ProtBERT model.

Each sequence was truncated to a maximum of 1024 length for training (out of 207 sequences, only 4 were truncated). The model, at both classification heads, performed classification at residue-level (token-level). Dataset was highly imbalanced; weighted Cross Entropy Loss function, given in eq. 1, was used by giving more weight to the positive (allosteric) class. Moreover, the dataset (Klausen et al., 2018) for secondary structure prediction task (secondary task) had 3 classes referring to (i) Alpha Helix (H), (ii) Beta Sheet (E), (iii) Coil (C), hence its head had 3 output units.

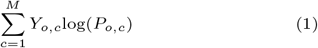

where *M* is total number of classes, *Y*_*o,c*_ is the ground truth and *P*_*o,c*_ is the predicted probability of observation *o* being in class *c*.

The pLM was fine-tuned in two settings: (a) **Base** - only allostery prediction head (no MTL) and (b) **MTL** - Allostery and Secondary Structure (SS) tasks comprising of two prediction heads. The architecture for finetuning the model, in MTL fashion, is given in Figure 2. No soft parameter sharing was used as the tasks are already similar. The model takes a protein sequence, and produces a 1024-*d* feature vector for each residue in a sequence of length *L*, hence giving a tensor of size *L* × 1024. Further, two heads, Secondary Structure (SS) classification head and Allostery Classification head, perform their own classification in order to fine-tune the model. Hence the **idea** was that, the multi-task learning handles *m* multiple learning tasks, denoted as 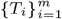, where the tasks share similarities but are not exactly the same. The goal is to improve the prediction performance of a model on a specific task *T*_*i*_ by leveraging information from other related tasks.

**Fig. 2.**
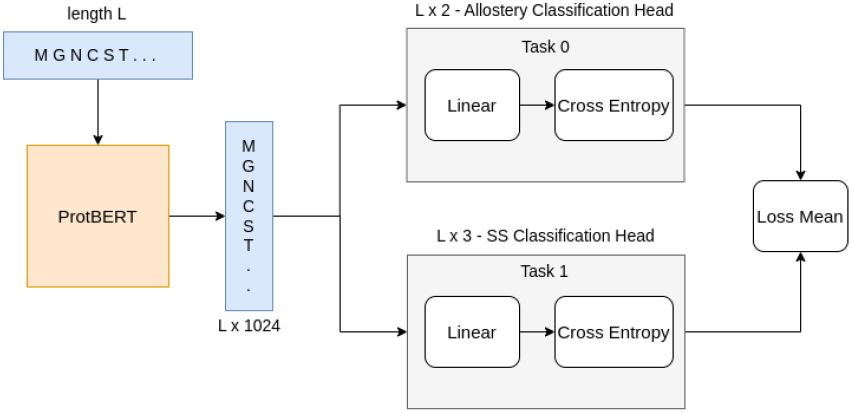
pLM finetuning framework of the proposed architecture. SS Head refers to Secondary Structure classification task. *L* is sequence length and for each residue (token) the model produces 1024*d* feature vector, hence *L ×* 1024. SS has 3 classes (*E* - Beta sheet, *H* - Alpha Helix, *C* - Coil) hence *L ×* 3 and allosteric head has two classes (positive or negative), hence *L ×* 2.

In this architecture (Figure 2), supervised learning is being performed and multi-task supervised learning setting means that each task in MTL is a supervised learning task, which models the functional mapping from data instances to labels. We are given *m* supervised tasks *T*_*i*_ for *i* = 1, …, *m*, each with its own training dataset 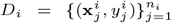 where 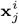 is a feature vector in *d*-dimensional space, and 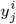 is the corresponding label (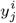 are discrete). For the *i*th task, there are *n*_*i*_ pairs of input data and labels. The aim of multi-task learning (MTL) is to learn *m* functions 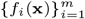 one for each task, such that each function 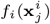 can accurately predict the associated label 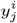. After these functions are trained, they can then be applied to new, unseen data points from the corresponding tasks to make predictions. Table 2 gives the hyperparameters that were used to fine-tune the pLM in MTL fashion.

**Table 2.**
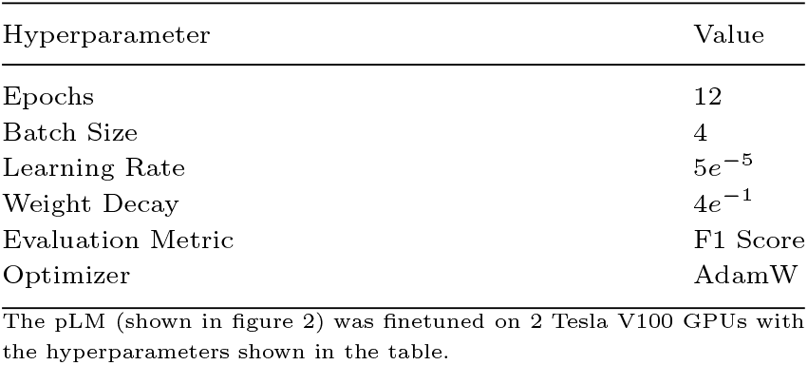
Hyperparamters for ProtBERT Finetuning (MTL)

#### 2.2.2. Pockets Extraction and Downstream Model Training

FPocket (Guilloux et al., 2009) was used to extract pockets from the protein structures. However, the scope of the work is to finetune the ProtBERT model in MTL setting and answer that, whether the finetuned pLM specifically in MTL fashion helps improve the prediction performance of allosteric pockets. For this, as language models cannot directly predict pockets/sites from a protein sequences, XGBoost and AutoML models were trained on combined FPocket features and features extracted from the finetuned pLM to predict whether a given pocket is an allosteric pocket or not.

#### 2.2.3. Training XGBoost

XGBoost was trained on pockets dataset. Each pocket’s features were concatenated with the pLM features such that only those pLM residue features were used for a certain pocket that were part of that pocket. It was trained by using binary logistic function as objective function with regularization *λ* = 0.15 and max tree depth of 7. To overcome the class imbalance problem, scale_pos_weight was set to the ratio of negative samples to the positive samples in training partition. The model was trained for 100 boosting rounds with 5-fold cross validation strategy, hence the scale_pos_weight parameter was calculated dynamically for each fold.

#### 2.2.4. Training AutoML

AutoML takes out all manual processing steps from preprocessing the dataset to model training and selection by automating this process into a pipeline. We used AutoGluon (Erickson et al., 2020) library for this purpose; it leverages multi-layer stacking with k-fold bagging. AutoGluon automatically selects the the layers and value of k during the training process. As mentioned in the Training XGBoost section 2.2.3, same dataset was used comprising of the finetuned pLM features and pocket features. We trained the AutoML models (pipeline) with default parameters on a single GPU, during which the pipeline trained several models with 3 layers of stacking and 8-folds.

## 3. Experimental Evaluation

As the dataset is highly imbalanced, the performance was evaluated using F1 score mainly but also keeping in mind Precision and Recall. F1 score is not only a measure of how many positives are found, but it also penalizes for the false positives that the method finds. Due to the high imbalance in the dataset, model achieved a high accuracy simply by predicting the majority class all the time, thus making the accuracy metric misleading. F1 score is more suitable in this case as it takes into account both precision (how many of the predicted positive instances are actually positive) and recall (how many of the actual positive instances are predicted positive). A model with a high F1 score is both good at avoiding false positives and false negatives, making it a more balanced measure than accuracy in imbalanced datasets.

## 4. Results and Discussion

### 4.1. Features’ Analysis

Feature exploration and analysis is an important step in ML. In this study, two types of features were used: FPocket features (representing pockets) and ProtBERT features (representing sequence residues). To analyze the class distribution, pockets features were concatenated with residue features (making up 1043-*d* feature vector corresponding to each pocket) and t-SNE (t-Distributed Stochastic Neighbour Embedding) (van der Maaten and Hinton, 2008) was performed to reduce dimensions from 1043 (1024+19) to 2 dimensions. Here *X* and *Y* are simply the names given to reduced dimensions.

Visually, the pockets do not cluster into distinct positive or negative partitions. However, it can be seen in Figure 3 that the majority of positive samples are shown in the upper right corner and top right of the plot. This shows that they share some similarities in high-dimensional space. The two clusters show some distinct protein classes but with some overlap in molecular weights. Top cluster mainly consists of molecular enzymes, and the top-right contains viral enzymes. Both clusters have similar average sizes (~110 kDa), but the top cluster includes extreme outliers (e.g., RNA polymerase holoenzyme at 867 kDa) and has a wider distance range (0.22 - 14.48 Å), however, the top-right cluster has lower median distance (3.87 Å), implying tighter spatial clustering of pockets. Top right cluster’s modulator types mostly consist of ATP, nucleotide analogs (e.g., cAMP), and small-molecule inhibitors (e.g., BPTES), however in the case of top-right one, it is mostly viral inhibitors (e.g., HIV RT inhibitor 7), antibiotics (e.g, ceftaroline), and kinase modulators. In addition, negative samples are scattered all over the plot, which could be a hint of a greater diversity within the negative class. Further detailed analysis is given in the supplementary material S2.1.

**Fig. 3.**
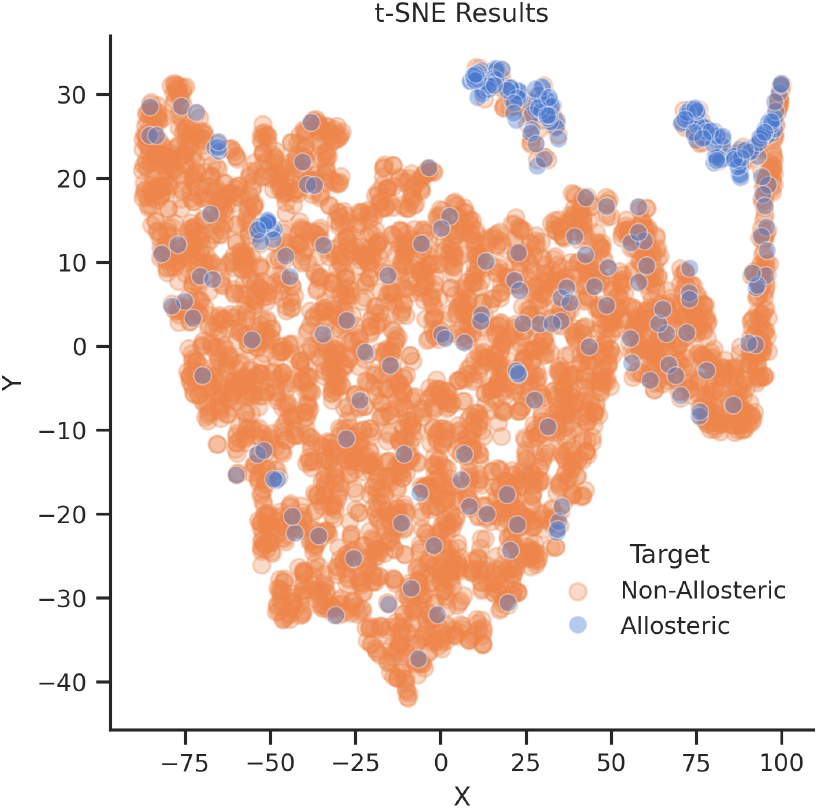
Allosteric and Non-Allosteric Pockets’ t-SNE Features Distribution

### 4.2. Results’ Analysis of Finetuning ProtBERT-BFD

To finetune the ProtBERT-BFD pLM, as mentioned in the 2.1 dataset section, each residue was labelled as either positive (allosteric) or negative (non-allosteric). BertTokenizer from HuggingFace was used to tokenize the sequences with sequence length restricted to 1024 size.

We finetuned the pLM in MTL setting and without MTL. Table 3 gives the results of models, where *MTL* refers to the pLM finetuned in MTL fashion and *Base* refers to without MTL. MTL-based model performs better than base model which proves that MTL increases the prediction performance. Moreover, we are also interested in higher precision (that means we are looking for accurate true positives), which MTL model gives us. Vig et al. (2021) interpreted attention in the ProtTrans models (Elnaggar et al., 2021) and they concluded that in the deeper layers of the LLM, heads give more attention to the binding sites and contacting residues, relatively to the earlier layers. Binding sites in proteins play an important role in interacting with other molecules that influence the protein’s functionality. Even with evolution, binding sites are conserved and remain unchanged as they have important role in protein (Kinjo and Nakamura, 2009). This is the reason that attention targets binding sites. Like the main binding sites, it can be conjectured that allosteric sites are also conserved to some extent. Hence, the model can give more attention to the conserved regions (allosteric pockets) even when other parts of the sequence may vary.

**Table 3.** ProtBERT Results’ Comparison - Base vs. MTL.

| Model | F1 | Precision | Recall |
| --- | --- | --- | --- |
| ProtBERT<sub>Base</sub> | 0.2688 | 0.2205 | **0.3442** |
| ProtBERT<sub>MTL</sub> | **0.2999** | **0.3193** | 0.2828 |

XGBoost and AutoML models were trained with FPocket only features providing us with baseline (Poc) results and we compare pLM based results with the baseline, given in tables 4 and 5.

**Table 4.** XGBoost Results’ Comparison - Base vs. MTL.

| Model (features) | F1 | Precision | Recall |
| --- | --- | --- | --- |
| Baseline (Poc) | 0.5586 | 0.5962 | 0.5254 |
| pLM<sub>Base</sub> | 0.7179 | 0.8077 | 0.6462 |
| pLM<sub>MTL</sub> | **0.8596** | **0.8909** | **0.8306** |
| pLM<sub>Base</sub> + Poc | 0.8361 | 0.8947 | 0.7846 |
| pLM<sub>MTL</sub> + Poc | **0.8870** | **0.9107** | **0.8644** |

**Table 5.** AutoML Results’ Comparison - Base vs. MTL.

| Model (features) | F1 | Precision | Recall |
| --- | --- | --- | --- |
| Baseline (Poc) | 0.5818 | 0.6275 | 0.5424 |
| pLM<sub>Base</sub> | 0.7500 | 0.8936 | 0.6462 |
| pLM<sub>MTL</sub> | **0.8649** | **0.9123** | **0.8136** |
| pLM<sub>Base</sub> + Poc | 0.8036 | 0.9036 | 0.6923 |
| pLM<sub>MTL</sub> + Poc | **0.8966** | **0.9231** | **0.8814** |

Additionally, we observed there were a small number of false negatives and positives prompting us to analyze it. The model struggled with complex and larger binding sites. Further details are given in the supplementary material (S2.2).

### 4.3. XGBoost Results’ Analysis

XGBoost was trained and tested on features making up of (a) only pLM-base features (*Base*), (b) only pLM-MTL features (*MTL*), (c) combination of pLM-base and FPocket features (*Base + Poc*), and (d) combination of pLM-MTL and FPocket features (*MTL + Poc*). Table 4 gives the results of XGBoost model (in the same order, a, b, c, and d) comparing results in groups of each two rows. In both cases, (with and without pocket features), features from MTL-based pLM outperform the model trained on features from base model. Highest F1 score achieved was 88.7%.

### 4.4. AutoML Results’ Analysis

AutoML was also trained with same settings as mentioned in the XGBoost section 4.3. Models trained on MTL-based pLM features outperformed than that of base pLM features. Highest F1 score achieved was 89.66%.

Models trained with MTL-pLM features have better recall and precision and overall they perform better than base pLM features based models.

### 4.5. Results Comparison and Discussion

The method discussed above, used pLM features and/or pocket features. Both results have been given with and without pocket features in order to show how pLM features affect the prediction performance of allosteric pockets. Results from XGBoost and AutoML are highly correlated; in fact we got a 0.9768 correlation score between the results of XGBoost and AutoML. Moreover, not only the pLM showed better performance but MTL-based pLM outperformed all other methods.

Allosteric pockets were ranked in different top positions namely, Top 1%, Top 3%, Top 5%, and Top 10%. Figure 4 shows that 90% of positive or predicted allosteric pockets by AutoML based on MTL-pLM features, are ranked in top 10% of the results.

**Fig. 4.**
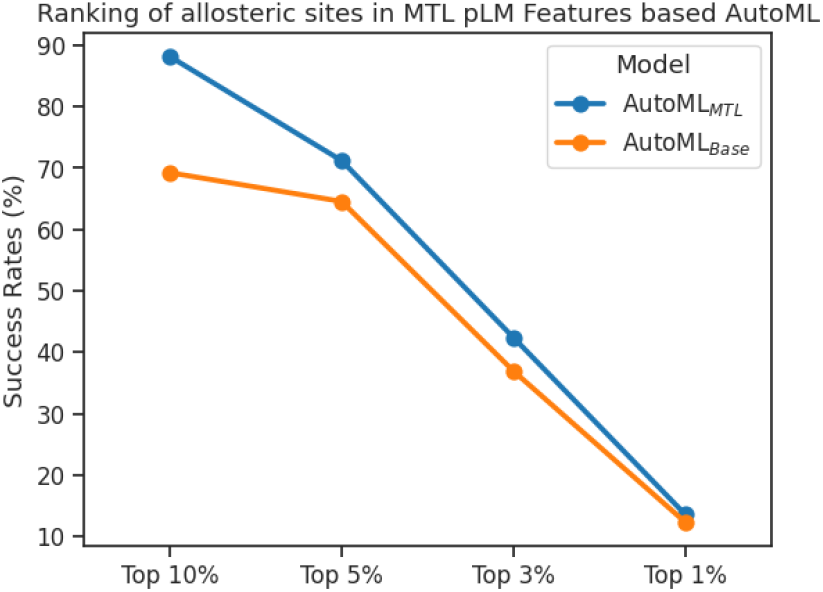
Ranking of Allosteric Pockets in MTL vs. base pLM Features based AutoML

Table 6 gives the overall results comparison of MTL-based models with previous approaches, with findings from previous approaches taken from their respective papers. Previous studies have used only pocket features to predict allosteric pockets while the models in this work have utilized the MTL-pLM features. As the model performs better with the combination of pLM_MTL_ and pocket features, the same model’s results are compared with previous approaches.

**Table 6.** Comparison of MTL-based pLM Models with Previous Models.

| Model (features) | F1 | Precision | Recall |
| --- | --- | --- | --- |
| SVM (Tian et al., 2023b) | 0.559 | 0.444 | 0.758 |
| XGBoost (Tian et al., 2023b) | 0.596 | 0.586 | 0.609 |
| LTR (Tian et al., 2023b) | 0.662 | 0.662 | 0.662 |
| AutoML (Tian et al., 2023a) | 0.701 | 0.850 | 0.616 |
| Ensemble (Tian et al., 2023a) | 0.782 | 0.726 | 0.847 |
| DeepAllo<sub>XGBoost</sub> | 0.887 | 0.911 | 0.864 |
| DeepAllo<sub>AutoML</sub> | **0.897** | **0.923** | **0.881** |

DeepAllo_*AutoML*_ model (trained on MTL based pLM features with pocket features) outperforms all previous models and has 12.8% increase over highest performing (Ensemble) (Tian et al., 2023a) model. Moreover, our model can predict a pocket with 90.5% confidence among the top 3 positions that is also higher than the Ensemble model’s top 3 i.e. 84.9%

## 5. Case Study

We tested our model on a case study protein (TOXIN B, PDB ID: 3PEE) that was not in our dataset. It is known to have allosteric pockets. AutoML_MTL_ model was used to predict the allosteric pockets. Top 3 pockets were selected by ordering predicted pockets in the descending order of probabilities.

In Figure 5, predicted pockets are marked in red, orange, and purple color. Red color shows that pocket with the highest probability, orange with the next highest probability, and the purple shows the third highest probability of an allosteric pocket. Moreover, the modulator is shown in green color. 1st pocket has 0.84 probability of being an allosteric pocket while remaining two pockets have 0.014 and 0.012 probability, respectively. The model was evaluated on 50% threshold i.e. a pocket must have atleast 0.5 probability to be predicted as allosteric, otherwise it would be considered negative.

**Fig. 5.**
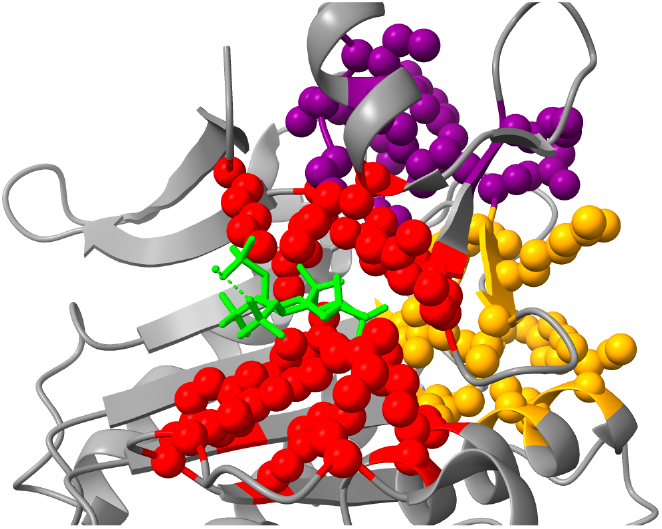
Predicted Allosteric Sites: Top 1st (Red), Top 2nd (Orange), Top 3rd (Purple), and Modulator (Green) - Atoms’ radius is scaled down for better spheres’ visualization.

The predicted pocket is the correct allosteric pocket. It can be observed that the predicted true allosteric pocket is near to the modulator (green). Moreover, third pocket (purple) is extremely far from the modulator, which is a proof that the model learns to predict correct allosteric pockets and takes into consideration the distance between the modulator and the allosteric pocket. It can be conjectured that the pLM_MTL_ features provide geometrical aspects of a protein and consequently help a model to differentiate between residues near the modulator and residues far from the modulator.

## 6. ProtBERT Explanation and Visualization

ProtBERT-BFD has 30 layers and 16 heads, resulting in a total of 30 × 16 = 480 distinct attention mechanisms. ProtBERT’s attention at different layers was also visualized in order to understand which head at which layer attends and which attention mechanism is followed. The visualization was performed for 5DKK protein. To keep it brief and make the point, neuron view for layer 7 and head 3 was captured, given in Figure 6. For the sake of brevity and visualization purposes, pocket residues were postfixed with an “underscore”. Attention is being visualized as lines connecting the residue being updated (left) with the residue being attended to. Color intensity reflects the attention weight; weights close to one are shown as dark lines, whereas weights close to zero appear as faint lines or not visible at all.

**Fig. 6.**
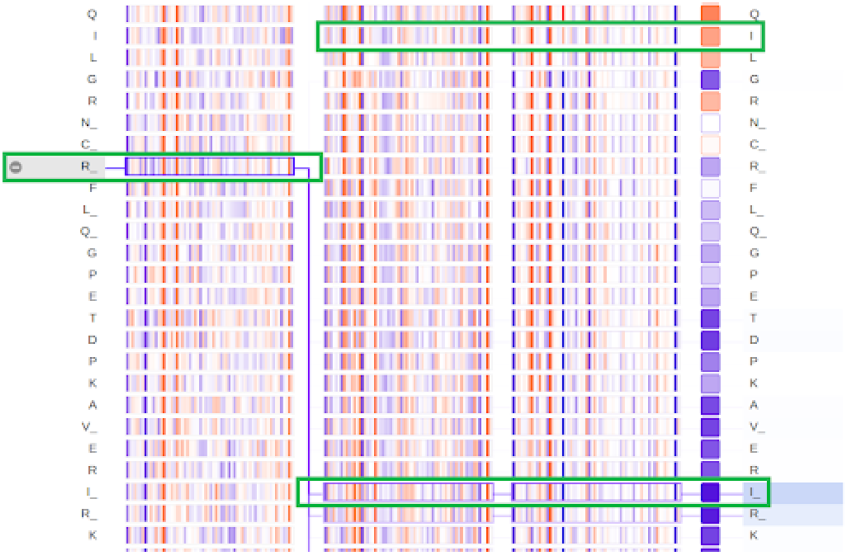
ProtBERT Neuron View: Layer 7 Head 3 - Three columns represent *Query, Key*, and *Value* in the attention matrix at this layer. Darker lines represent higher attention values; the column with square boxes has negative (orange) and positive (blue) attention values. Highlighted residues (with green rectangle) have same structural distance from R, however model learns to attend to allosteric residue (represented with an underscore).

Figure 6 gives the neuron view at layer 7 and head 3. First three columns represent the *Query, Key*, and *Value* in the Attention mechanism. Residues in question are highlighted with green rectangles. Residue *R* has same Euclidean distance of 10.5 Å from both residues *I* and *I*, however, it gives attention to *I* residue and does not attend to residue *I* (non-allosteric residue). Although, the geometrical distance is the same, it can capture the allosteric pocket residues even though, in the sequence, *I* is at a long distance from *R* compared to the distance from *I*.

The pLM mostly focuses on delimiter tokens in the initial layers and as the input passes through deeper layers, the model can make better connections among the residues (tokens). Several other patterns also make their presence known in the model such as attending to previous residue in the sequence and attending to next residue in the sequence.

## 7. Limitations

The language model can predict allosteric residues but it cannot predict allosteric pockets on its own. Moreover, the F1 score of the pLM is not that high and it might give false positives (as a standalone version). Secondly, the results of XGBoost and AutoML models are derived predictions that come from FPocket’s detected pockets. Whole architecture is more of an ensemble model that depends on FPocket for pockets’ extraction. Due to the unavailability of huge dataset and no access to experimental methods, it is hard to test the model for new allosteric proteins (this stands for all previous computational approaches in the literature). Moreover, the pLM exhibits a lot of attention mechanism and sometimes they are irrelevant; attending to CLS tokens, sometimes, may render the results incorrect. Last but not least, the pLM gives more attention to conserved sites, although allosteric sites are little bit conserved but they are not as conserved in protein families as the orthosteric sites are, making it hard for the pLM to make connections.

## 8. Conclusion

This work was aimed to leverage finetuned pLM specifically in Multitask Learning fashion in order to see if the prediction performance of allosteric pockets, over current approaches in the literature, improves or not. As in the published literature, ML and NMA approaches have been utilized to predict allosteric pockets in proteins; pLMs have never been utilized in this domain of study. Extending base pLM for this task, Multitask Learning was used to improve the prediction performance of allosteric pockets, by using secondary structure prediction as secondary task.

ProtBERT-BFD was finetuned and used as feature extractor for XGBoost and AutoML models. FPocket was used to extract pockets from allosteric proteins that was used as input for XGBoost and AutoML. The pLM features combined with pocket features were fed into XGBoost and AutoML, that resulted in classifying pockets as allosteric or non-allosteric. Due to high imbalance in the dataset F1 score was chosen as evaluation metric and the proposed model achieved 89.66% F1 score.

Additionally, a case study was performed to see the top 3 pockets and the model predicted the correct allosteric pocket as top 1 position with 99% confidence, proving that pLM finetuned with MTL do improve prediction performance of allosteric pockets in proteins.

Interpretation and demystification of a deep learning (DL) model is an important aspect. An effort was made to explain the pLM and how it captures the allosteric residues. Several attention mechanisms were identified. Visual interpretation from these patterns shows distinct information, however, the perceived information could be subjective.

In future, leveraging more sophisticated pLM models such as ProtT5 (based on T5 architecture) is expected to further improve the prediction performance of allosteric pockets.

## Data Availability Statement

The source code is available on https://github.com/ku-cosbi/deepallo.

## Author contributions statement

M.K., O.K., and A.G. conceived the study. M.K. conducted the experiments and analysed the results. M.K., O.K., and A.G. wrote and reviewed the manuscript.

## Acknowledgements

This work is supported by the Koç University İş Bank - Artificial Intelligence (KUIS AI) center and the Scientific and Technological Research Council of Turkey (TÜBİTAK) project 120C120. We would like to thank the people at COSBI lab for their invaluable support and to the IT department for managing the KUACC server, enabling us to run our huge models with GPU availability.

## Footnotes

1 https://mdl.shsmu.edu.cn/ASD/

2 https://github.com/smu-tao-group/PASSerRank

