## Supplementary Material for "DeepAllo: Allosteric Site Prediction using Protein Language Model (pLM) with Multitask Learning"

### DeepAllo: Supplementary Material

#### S1. Proteins Dataset with Names and PDB IDs

Table S1: List of 207 proteins and their PDB IDs in alphabetical order.

| No. | Protein Name | PDB ID |
|---|---|---|
| 1 | Protein (Bovine Seminal Ribonuclease) | 11BG |
| 2 | Carbamoyl Phosphate Synthetase (Large Chain) | 1A9X |
| 3 | Glutamine Phosphoribosylpyrophosphate Amidotransferase | 1AO0 |
| 4 | Alpha-Amylase | 1AQM |
| 5 | Protein (5-Aminolevulinic Acid Dehydratase) | 1B4K |
| 6 | Protein (Purine Repressor) | 1BDH |
| 7 | Tryptophan Synthase | 1BEU |
| 8 | Serine Hydroxymethyltransferase, Cytosolic | 1BJ4 |
| 9 | Phosphoribosyl Pyrophosphate Synthetase | 1DKR |
| 10 | Atrial Natriuretic Peptide Receptor A | 1DP4 |
| 11 | Lac Repressor | 1EFA |
| 12 | Cytochrome P450Eryf | 1EUP |
| 13 | Rad50 Abc-Atpase | 1F2U |
| 14 | O-Acetylserine Sulfhydrylase | 1FCJ |
| 15 | Phosphoenolpyruvate Carboxylase | 1FIY |
| 16 | Receptor-Type Adenylate Cyclase Gresag 4.1 | 1FX2 |
| 17 | Hemoglobin Alpha Chain | 1G9V |
| 18 | Annexin V | 1HAK |
| 19 | Anaerobic Ribonucleotide-Triphosphate Reductase | 1HK8 |
| 20 | S-Adenosylmethionine Decarboxylase Beta Chain | 1I72 |
| 21 | Anthranilate Synthase | 1I7S |
| 22 | 3-Deoxy-D-Arabino-Heptulosonate-7-Phosphatesynthase | 1KFL |
| 23 | L-Lactate Dehydrogenase | 1LDN |
| 24 | Tryptophan-Trna Ligase | 1M83 |
| 25 | Sulfate Adenylyltransferase | 1M8P |
| 26 | Cyclic Nucleotide Phosphodiesterase 2A | 1MC0 |
| 27 | S-Adenosylmethionine Decarboxylase Proenzyme | 1MSV |
| 28 | Fk506-Binding Protein | 1NSG |
| 29 | Amine Oxidase [Flavin-Containing] B | 1OJ9 |
| 30 | Ribonucleoside-Diphosphate Reductase 2 Alphachain | 1PEQ |
| 31 | Nad-Dependent Malic Enzyme, Mitochondrial | 1PJ2 |
| 32 | Protein (Hexokinase) | 1QHA |
| 33 | Parathion Hydrolase | 1QW7 |
| 34 | Repressor Protein | 1R1V |
| 35 | Integrin Alpha-L | 1RD4 |
| 36 | Caspase-7 | 1SHL |
| 37 | Protein-Tyrosine Phosphatase, Non-Receptor Type 1 | 1T48 |
| 38 | Cgmp-Specific 3',5'-Cyclic Phosphodiesterase | 1TBF |
| 39 | Atp-Dependent Clp Protease Atp-Binding Subunit Clpx | 1UM8 |
| 40 | B-Raf Proto-Oncogene Serine/Threonine-Protein Kinase | 1UWH |
| 41 | Glyceraldehyde-3-Phosphate Dehydrogenase (Nadp+) | 1UXN |
| 42 | Beta-Lactamase Shv-1 | 1VM1 |
| 43 | Uracil Phosphoribosyltransferase | 1VST |
| 44 | Cytochrome P450 3A4 | 1W0F |
| 45 | Acetyl-Coenzyme A Carboxylase | 1W96 |
| 46 | Hut Operon Positive Regulatory Protein | 1WPU |
| 47 | Ribonucleotide Reductase, B12-Dependent | 1XJE |
| 48 | Reca Protein | 1XMV |

Continued on next page

Table S1 – continued from previous page

| No. | Protein Name | PDB ID |
|---|---|---|
| 49 | Arginine Repressor | 1XXB |
| 50 | Coagulation Factor VII | 1YGC |
| 51 | Allantoate Amidohydrolase | 1Z2L |
| 52 | Copper-Containing Nitrite Reductase | 1ZDS |
| 53 | Dna-Directed Rna Polymerase Alpha Chain | 2A69 |
| 54 | Divalent Cation Transport-Related Protein | 2BBH |
| 55 | Acetylglutamate Kinase | 2BTY |
| 56 | Pyruvate Dehydrogensae Kinase Isoenzyme 2 | 2BU2 |
| 57 | Aspartokinase | 2CDQ |
| 58 | Protein (Arginase) | 2CEV |
| 59 | Protein-Arginine Deiminase Type Iv | 2DEW |
| 60 | 235Aa Long Hypothetical Biotin-[Acetyl-Coa-Carboxylase]Ligase | 2DJZ |
| 61 | Bira Bifunctional Protein | 2EWN |
| 62 | Dna Repair And Recombination Protein Rada | 2FPL |
| 63 | Estrogen Receptor Beta | 2FSZ |
| 64 | Deoxycytidylate Deaminase | 2HVW |
| 65 | D-Alanine-D-Alanine Ligase | 2I80 |
| 66 | Acetylcholinesterase | 2J3Q |
| 67 | Glutamate Racemase | 2JFN |
| 68 | Glutamate Racemase | 2JFZ |
| 69 | Myosin-2 Heavy Chain | 2JHR |
| 70 | DNA Mismatch Repair Protein MSH2 | 2O8D |
| 71 | Thymidylate Synthetase | 2ONB |
| 72 | Nitric Oxide Synthase, Inducible | 2ORO |
| 73 | Virulence Factor Regulator | 2OZ6 |
| 74 | L-Asparaginase I | 2P2N |
| 75 | Arginine Repressor | 2P5M |
| 76 | D-3-Phosphoglycerate Dehydrogenase | 2PA3 |
| 77 | Kinesin-Like Protein Kif11 | 2PG2 |
| 78 | Isomerase Domain Of Glutamine-Fructose-6-Phosphatetransaminase (Isomerizing) | 2POC |
| 79 | Hyperpolarization-Activated (Ih) Channel | 2PTM |
| 80 | Phenylpyruvate Decarboxylase | 2Q5Q |
| 81 | Transporter | 2Q6H |
| 82 | Fructose-Bisphosphatase | 2Q8M |
| 83 | Caspase-7 | 2QLJ |
| 84 | Prephenate Dehydratase | 2QMX |
| 85 | Uncharacterized Protein Vca0042 | 2RDE |
| 86 | Amp-Activated Protein Kinase Catalytic Subunit Alpha-1 | 2V8Q |
| 87 | Atp Phosphoribosyltransferase | 2VD3 |
| 88 | Bifunctional Protein Glmu | 2VD4 |
| 89 | Pyruvate Decarboxylase Isozyme 1 | 2VK1 |
| 90 | Tetracycline Resistance Repressor Protein | 2VPR |
| 91 | Ribonucleoside-Diphosphate Reductase Largesubunit | 2WGH |
| 92 | Glucosamine-6-Phosphate Deaminase | 2WU1 |
| 93 | 2-Oxoglutarate Decarboxylase | 2Y0P |
| 94 | Nickel And Cobalt Resistance Protein Cnrr | 2Y39 |
| 95 | 2-C-Methyl-D-Erythritol 4-Phosphate Cytidylyltransferase,Chloroplastic | 2YC3 |
| 96 | Androgen Receptor | 2YLO |
| 97 | Glycogen Phosphorylase, Liver Form | 2ZB2 |
| 98 | Putative Uncharacterized Protein Ph0207 | 3ATH |
| 99 | Glucose-1-Phosphate Adenylyltransferase | 3BRK |
| 100 | Glutamate Receptor, Ionotropic Kainate 1 | 3C35 |
| 101 | Protein RECA | 3CMW |
| 102 | Transthyretin | 3CN0 |
| 103 | Heat Shock Protein Homolog Sse1 | 3D2E |
| 104 | Putative Acetylglutamate Synthase | 3D2P |
| 105 | Cone Cgmp-Specific 3',5'-Cyclic Phosphodiesterase Subunitalpha' | 3DBA |
| 106 | Carbonic Anhydrase 2 | 3E2A |
| 107 | Cyclic Nucleotide-Binding Protein | 3E5U |

Continued on next page

Table S1 – continued from previous page

| No. | Protein Name | PDB ID |
|---|---|---|
| 108 | Dual Specificity Mitogen-Activated Protein Kinase Kinase 1 | 3EQC |
| 109 | Glud1 Protein | 3ETD |
| 110 | Endothelial Pas Domain-Containing Protein 1 | 3F1O |
| 111 | Alpha-Isopropylmalate Synthase | 3F6H |
| 112 | Leukotriene A-4 Hydrolase | 3FUD |
| 113 | Rtx Toxin Rtxa | 3GCD |
| 114 | Rna-Directed Rna Polymerase | 3GNV |
| 115 | Casein Kinase Ii Subunit Alpha | 3H30 |
| 116 | Exodeoxyribonuclease I | 3HL8 |
| 117 | Protein Fimx | 3HV8 |
| 118 | Reverse Transcriptase/Ribonuclease H | 3I0S |
| 119 | Transcriptional Regulator, Crp/Fnr Family | 3I54 |
| 120 | Camp-Specific 3',5'-Cyclic Phosphodiesterase 4D | 3IAD |
| 121 | Macrophage Migration Inhibitory Factor | 3IJG |
| 122 | Serine/Threonine-Protein Kinase Chk1 | 3JVS |
| 123 | Peroxisome Proliferator-Activated Receptor Gamma | 3K8S |
| 124 | Protease | 3KF0 |
| 125 | Probable 3-Deoxy-D-Arabino-Heptulosonate 7-Phosphatesynthase Arog | 3KGF |
| 126 | Protein Mj1225 | 3KH5 |
| 127 | Caspase-3 | 3KJF |
| 128 | Aspartokinase | 3L76 |
| 129 | Global Nitrogen Regulator | 3LA3 |
| 130 | Arginine Repressor | 3LAP |
| 131 | Isocitrate Dehydrogenase Kinase/Phosphatase | 3LCB |
| 132 | Serum Albumin | 3LU6 |
| 133 | Pantothenate Kinase 3 | 3MK6 |
| 134 | Suppressor Of Kinetochore Protein 1 | 3MKS |
| 135 | Prephenate Dehydratase | 3MWB |
| 136 | Farnesyl Pyrophosphate Synthase | 3N1V |
| 137 | Orf 17 | 3NJQ |
| 138 | Mitogen-Activated Protein Kinase 8 | 3O2M |
| 139 | Camp-Dependent Protein Kinase Regulatory Subunit | 3OF1 |
| 140 | 3-Phosphoinositide-Dependent Protein Kinase 1 | 3OTU |
| 141 | Myosin Heavy Chain Kinase A | 3PDT |
| 142 | Toxin B | 3PEE |
| 143 | Phospho-2-Dehydro-3-Deoxyheptonate Aldolase | 3PG9 |
| 144 | Udp-Glucose 6-Dehydrogenase | 3PJG |
| 145 | Camp-Dependent Protein Kinase Type I-Alpha Regulatorysubunit | 3PNA |
| 146 | Udp-Glucose 6-Dehydrogenase | 3PTZ |
| 147 | Adenosine Receptor A2A,Lysozyme Chimera | 3QAK |
| 148 | Nmda Glutamate Receptor Subunit | 3QEM |
| 149 | Ribonucleotide Reductase R1 Protein | 3R1R |
| 150 | Ribonucleoside-Diphosphate Reductase Large Chain 1 | 3RSR |
| 151 | Dihydrodipicolinate Synthase | 3TCE |
| 152 | Probable Dna Double-Strand Break Repair Rad50 Atpase | 3THO |
| 153 | Insulin-Degrading Enzyme | 3TUV |
| 154 | 6-Phosphofructokinase Isozyme 2 | 3UMO |
| 155 | Glutaminase Kidney Isoform, Mitochondrial | 3UO9 |
| 156 | Glutaminase Kidney Isoform, Mitochondrial | 3VOZ |
| 157 | Penicillin Binding Protein 2 Prime | 3ZG0 |
| 158 | Nickel And Cobalt Resistance Protein Cnrr | 3ZG1 |
| 159 | Glucose-1-Phosphate Thymidylyltransferase | 3ZLK |
| 160 | Choline Kinase Alpha | 3ZM9 |
| 161 | Uridylate Kinase | 4A7W |
| 162 | Lysine Acetyltransferase | 4AVC |
| 163 | Non-Structural Protein 4A, Serine Protease Ns3 | 4B6E |
| 164 | 3-Oxoacyl-[Acyl-Carrier-Protein] Reductase Fabg | 4BNU |
| 165 | Udp-N-Acetylglucosamine Pyrophosphorylase | 4BQH |
| 166 | Nad-Dependent Protein Deacetylase Sirtuin-3,Mitochondrial | 4C7B |

Continued on next page

Table S1 – continued from previous page

| No. | Protein Name | PDB ID |
|---|---|---|
| 167 | Integrase | 4CIG |
| 168 | Adenylate Cyclase Type 10 | 4CLL |
| 169 | Glucokinase | 4DCH |
| 170 | Camp-Dependent Protein Kinase Catalytic Subunit Alpha | 4DFZ |
| 171 | Rac-Alpha Serine/Threonine-Protein Kinase | 4EJN |
| 172 | Peld | 4EU0 |
| 173 | Neurolysin, Mitochondrial | 4FXY |
| 174 | Aspartate Carbamoyltransferase Catalytic Chain | 4FYV |
| 175 | Plasminogen Activator Inhibitor 1 | 4G8O |
| 176 | Dihydrodipicolinate Synthase | 4HNN |
| 177 | Prmt3 Protein | 4HSG |
| 178 | Pyruvate Kinase 1 | 4HYW |
| 179 | Magnesium Transport Protein Cora | 4I0U |
| 180 | Mucosa-Associated Lymphoid Tissue Lymphoma Translocationprotein 1 | 4I1R |
| 181 | Heat Shock 70Kda Protein 1A Variant | 4IO8 |
| 182 | Isocitrate Dehydrogenase [Nadp], Mitochondrial | 4JA8 |
| 183 | Citrate Synthase | 4JAF |
| 184 | Chaperone Protein Dnak | 4JN4 |
| 185 | N-Acetylglutamate Kinase / N-Acetylglutamate Synthase | 4KZT |
| 186 | Gtpase Kras | 4LUC |
| 187 | Copper-Sensitive Operon Repressor (Csor) | 4M1P |
| 188 | Chimera Protein Of C-C Chemokine Receptor Type 5 Andrubredoxin | 4MBS |
| 189 | Ubiquitin-Conjugating Enzyme E2 R1 | 4MDK |
| 190 | Dna Double-Strand Break Repair Rad50 Atpase | 4NCJ |
| 191 | Udp-N-Acetylglucosamine 2-Epimerase | 4NES |
| 192 | Dihydropteroate Synthase | 4NIR |
| 193 | Cytosolic Imp-Gmp Specific 5'-Nucleotidase | 4OHF |
| 194 | Metabotropic Glutamate Receptor 5, Lysozyme, Metabotropicglutamate Receptor 5 Chimera | 4OO9 |
| 195 | Bira Bifunctional Protein | 4OP0 |
| 196 | Soluble Cytochrome B562, Metabotropic Glutamate Receptor 1 | 4OR2 |
| 197 | Deoxycytidylate Deaminase | 4P9D |
| 198 | Phosphofructokinase | 4PFK |
| 199 | Vitamin D3 Receptor A | 4Q0A |
| 200 | Isoprenyl Transferase | 4Q9O |
| 201 | Deoxynucleoside Triphosphate Triphosphohydrolase Samhd1 | 4QFX |
| 202 | E3 Ubiquitin-Protein Ligase Rnf146 | 4QPL |
| 203 | Pyruvate Carboxylase | 4QSK |
| 204 | Prostaglandin G/H Synthase 2 | 4RUT |
| 205 | Eukaryotic Translation Initiation Factor 4E | 4TPW |
| 206 | Cystathionine Beta-Synthase | 4UUU |
| 207 | Chorismate Mutase | 5CSM |

#### S2. Data and Results' Analysis

##### S2.1. t-SNE Distribution of Pocket Features

The features of the t-SNE pocket were further analyzed to see their classes, weights, and distance distributions. Allosteric pockets are mainly close to the modulator and have an average size of ∼110 kDa. Refer to the figures S1, S2, and S3.

##### S2.2. Model Analysis on Confusion Matrix - FP vs. FN

False negatives share some similarities with true positives i.e., large volumes ($3754.79 \pm 3394.02$ Å$^3$) and high residue counts ($16.43 \pm 9.52$), which indicates that the model struggles with complex, larger binding sites. False positives show intermediate characteristics, with volumes ($2878.87 \pm 2990.05$ Å$^3$) and residue counts ($13.80 \pm 10.38$) falling between typical binding and non-binding pockets. This pattern is supported by substantial effect sizes between false positives and true negatives for both volume (Cohen's $d$ = 0.77) and number of residues (Cohen's $d$ = 1.14). Hydrophobicity shows minimal variation across all categories (p=0.8891 in Kruskal-Wallis test), showing that it is not a decisive factor in the model's predictions. The model appears to struggle most with "borderline" cases where size and volume metrics fall between typical binding and non-binding ranges, and with complex pockets having higher residue counts. While the small number of false predictions indicates good overall performance, these edge cases where multiple features do not clearly align with typical characteristics represent the model's main weakness. The correlation

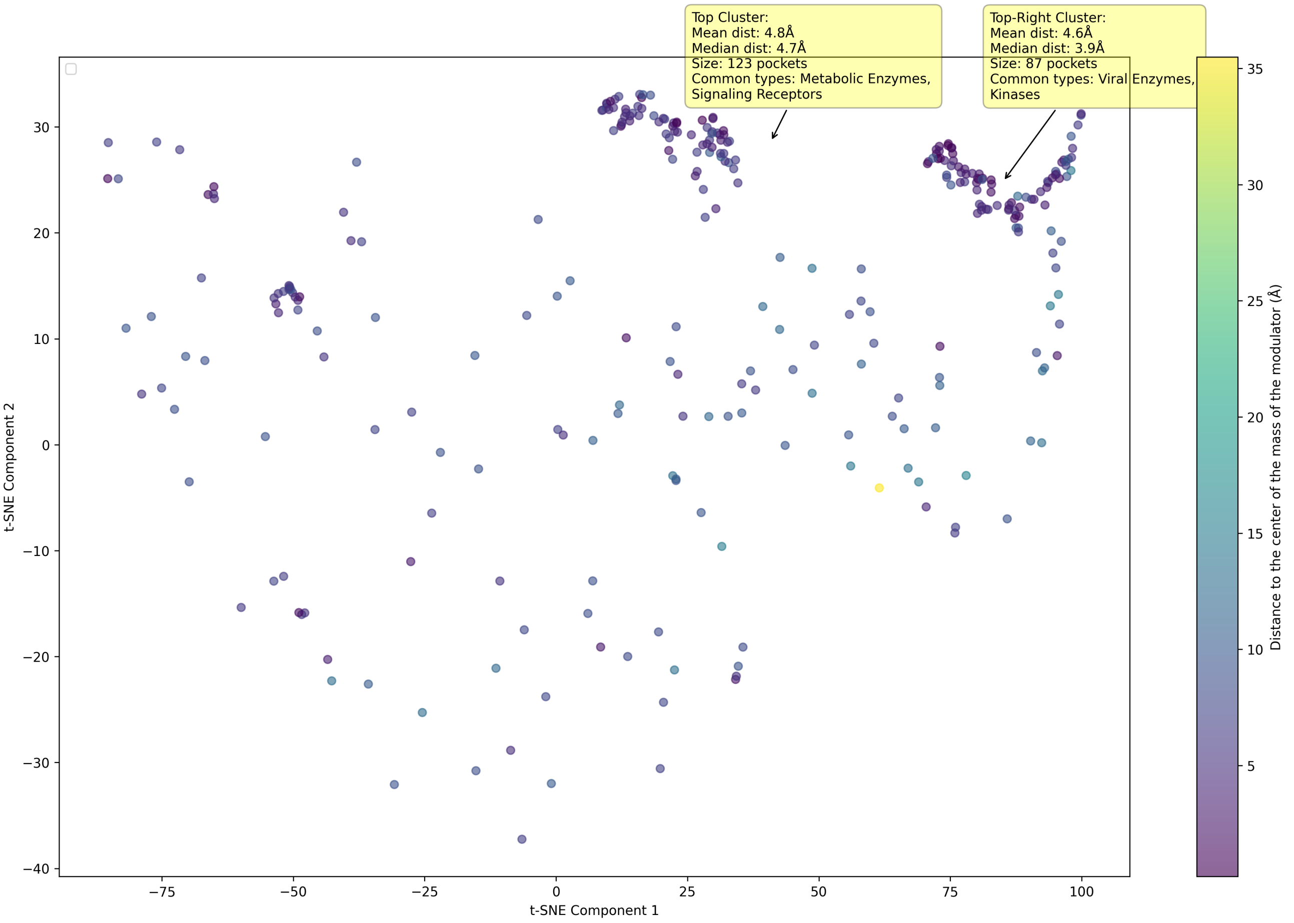


**Fig. S1.** Top and Top-right Clusters of Allosteric Pockets - Axes' labels are random names that represent the two features given by t-SNE analysis.

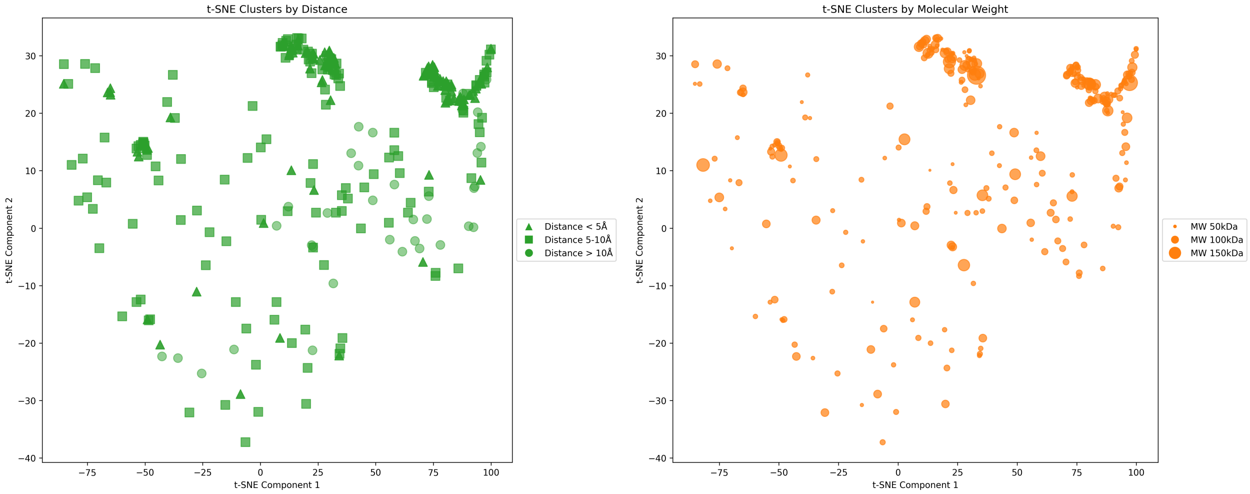


**Fig. S2.** Allosteric Pocket Clusters - Average mean weight (MW) for allosteric pockets is  110 kDa, however, the top-right cluster has a lower median distance (3.87 Å) between the center of mass of a pocket and the center of mass of its modulator compared to the top cluster (4.65 Å). (Axes' labels are random names that represent the two features given by t-SNE analysis.)

matrix shows hydrophobicity is not highly correlated with other metrics (*size*, *volume*, and *number of residues*). These trends are given in figures S4, and S5.

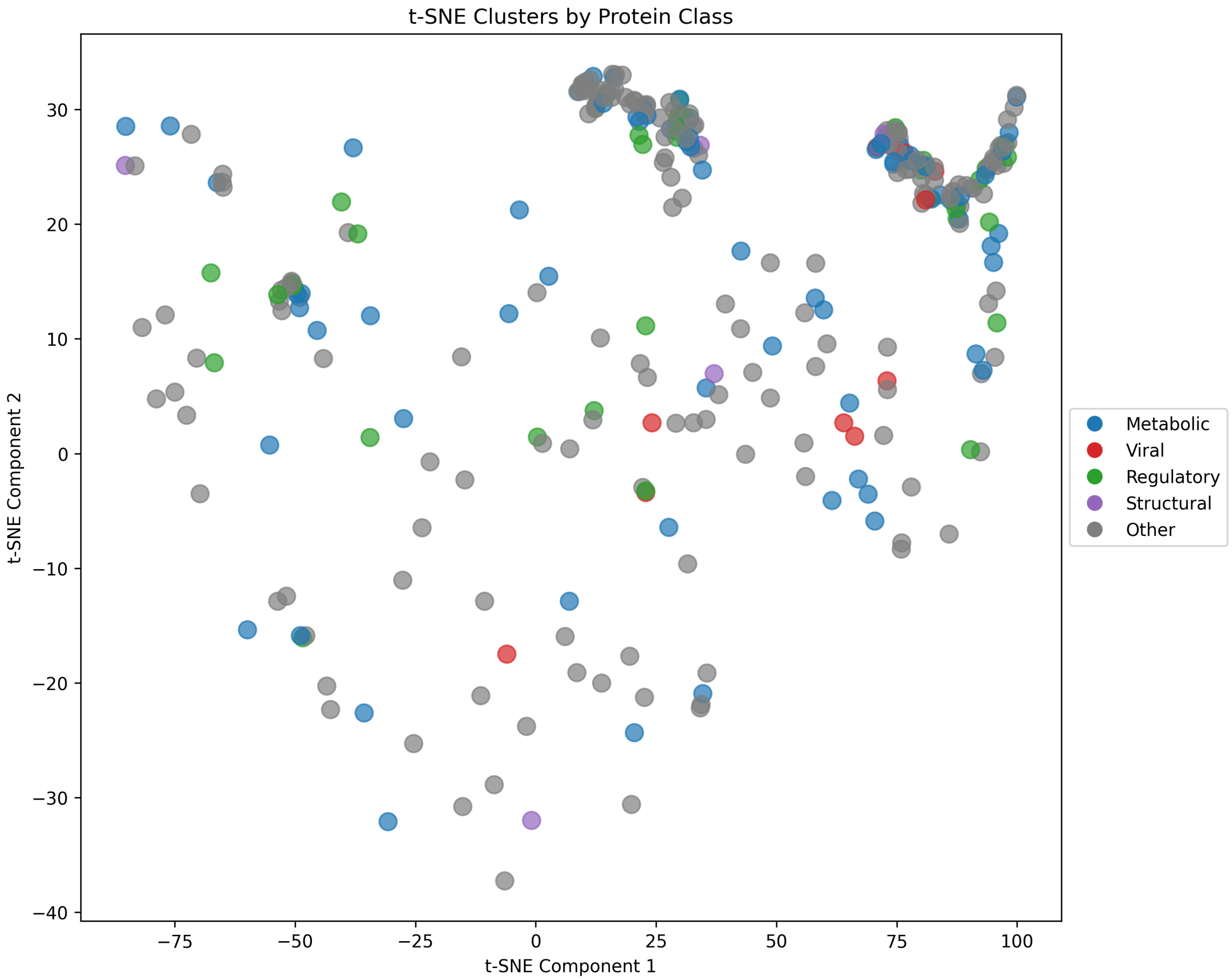


**Fig. S3.** Allosteric Pocket Clusters with Protein Classifications - Top right cluster's modulator types mostly consist of ATP, nucleotide analogs (e.g., cAMP), and small-molecule inhibitors (e.g., BPTES), however in the case of top-right one, it is mostly viral inhibitors (e.g., HIV RT inhibitor 7), antibiotics (e.g, ceftaroline), and kinase modulators. (Axes' labels are random names that represent the two features given by t-SNE analysis.)

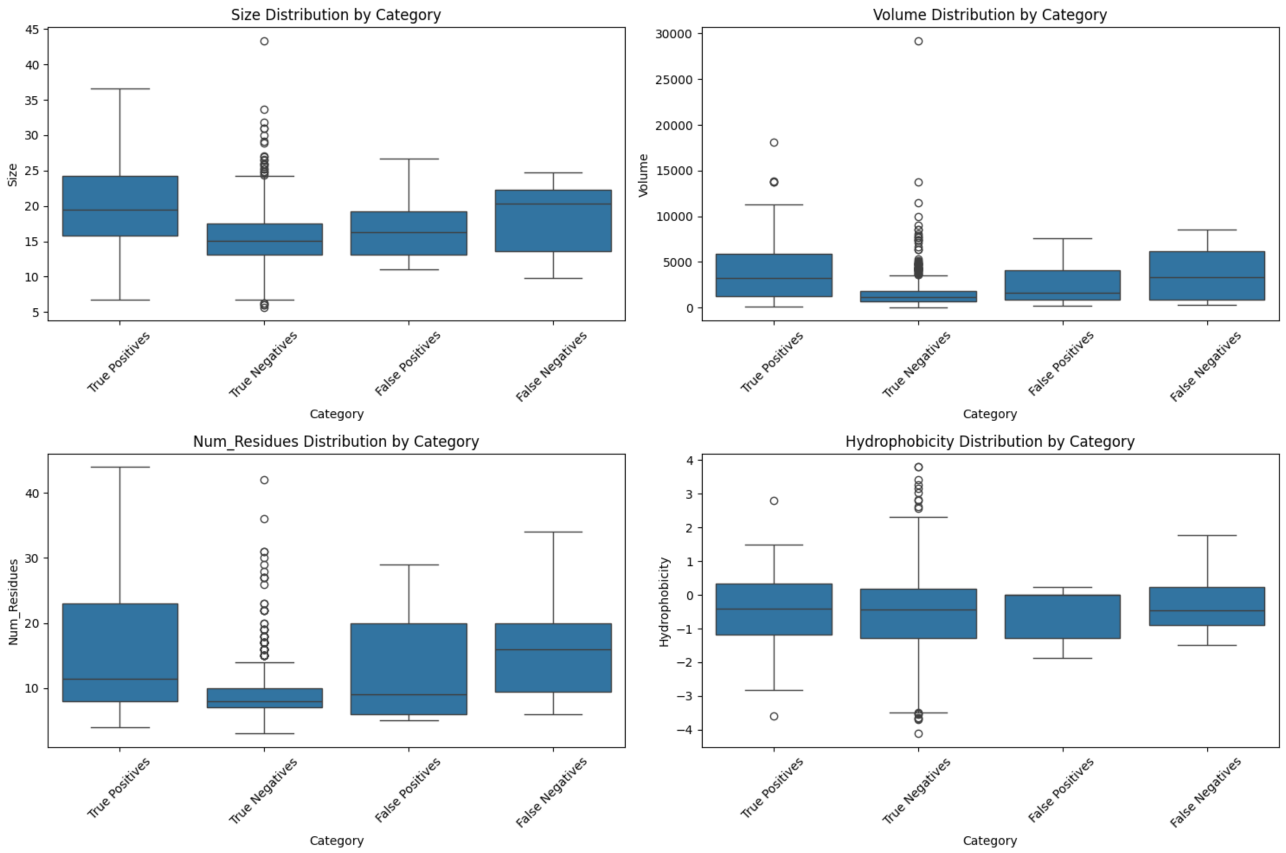


**Fig. S4.** Pockets Distribution by Size, Volume, Hydrophobicity, and Number of Residues - Size represents the maximum pairwise distance between any two points in the pocket (in Å). Volume is calculated as the product of the bounding box dimensions containing all pocket points (in Å$^3$). Num_Residues indicates the count of amino acid residues lining the pocket.

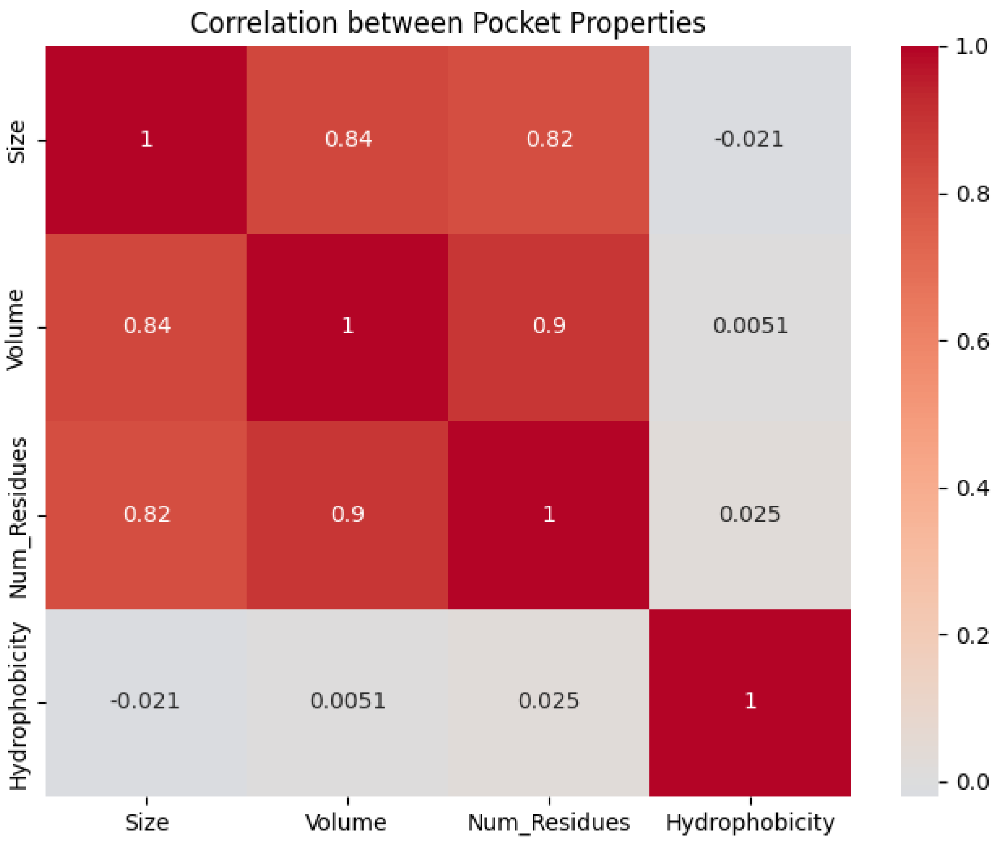


**Fig. S5.** Correlation heatmap showing the relationships between different pocket properties. Strong positive correlations are observed between Size, Volume, and Number of Residues (0.82-0.9), indicating these geometric properties are highly interrelated. In contrast, Hydrophobicity shows negligible correlations with all other properties (correlation coefficients between -0.021 and 0.025), suggesting that the chemical nature of the pocket is independent of its physical dimensions.
